# Generation and Maturation of Human iPSC-derived Cardiac Organoids in Long Term Culture

**DOI:** 10.1101/2022.03.07.483273

**Authors:** Ece Ergir, Jorge Oliver-De La Cruz, Soraia Fernandes, Marco Cassani, Francesco Niro, Daniel Sousa, Jan Vrbský, Vladimír Vinarský, Ana Rubina Perestrelo, Doriana Debellis, Francesca Cavalieri, Stefania Pagliari, Heinz Redl, Peter Ertl, Giancarlo Forte

**Author notes:** **Corresponding author:** Giancarlo Forte, PhD, Center for Translational Medicine (CTM), International Clinical Research Center (ICRC), St. Anne’s University Hospital, Studentska 6, Brno, Czech Republic, 62500.

## Abstract

Cardiovascular diseases remain the leading cause of death worldwide; hence there is an increasing focus on developing physiologically relevant *in vitro* cardiovascular tissue models suitable for studying personalized medicine and pre-clinical tests. Despite recent advances, models that reproduce both tissue complexity and maturation are still limited.

We have established a scaffold-free protocol to generate multicellular, beating and self-organized human cardiac organoids (hCO) *in vitro* from hiPSCs that can be cultured for long term. This is achieved by differentiation of hiPSC in 2D monolayer culture towards cardiovascular lineage, followed by further aggregation on low-attachment culture dishes in 3D. The generated human cardiac organoids (hCOs) containing multiple cell types that physiologically compose the heart, gradually self-organize and beat without external stimuli for more than 50 days. We have shown that 3D hCOs display improved cardiac specification, survival and maturation as compared to standard monolayer cardiac differentiation. We also confirmed the functionality of hCOs by their response to cardioactive drugs in long term culture. Furthermore, we demonstrated that hCOs can be used to study chemotherapy-induced cardiotoxicity.

This study could help to develop more physiologically-relevant cardiac tissue models, and represent a powerful platform for future translational research in cardiovascular biology.

## Introduction

Cardiovascular diseases (CVD) remain the leading cause of death worldwide^1–4^, and developing new therapies is still a major challenge, since a significant number of drug candidates fail to pass clinical trials, or are withdrawn from the market due to adverse effects^5–7^. In order to approve safer and more effective therapies, there is an increasing demand to develop faithful models of human heart tissue for pre-clinical research^7^. While recent technologies provide some insight into how human CVDs can be modelled *in vitro*, a comprehensive overview of the complexity of the human heart remains elusive due to the limited cellular heterogeneity, physiological complexity, or maturity of the constructs produced^8^. Furthermore, animal models may not always faithfully reflect the unique features of human biology and disease, and could give rise to ethical concerns^9–11^.

Induced pluripotent stem cell (iPSC) technology has been revolutionary for the differentiation and derivation of cardiomyocytes for personalized disease modelling and drug testing^8,12–20^. However, when cultured in 2D as models for development, disease and toxicology^18,21,22^, cardiomyocytes do not reflect the 3D complexity of the native tissue, where the geometry, the presence of different cell types and their interaction with the extracellular matrix (ECM) play a key role.

Early 3D cardiac tissue models called “cardiospheres” were developed by culturing human heart tissue biopsies, or by mixing non-isogenic populations of cardiomyocytes, non-myocytes and biocompatible hydrogels^23–25^; however, such microtissues generally had limited culture continuity, self-organization and failed to capture the heterogeneity characteristic of organotypic models. Recently, cardiac tissue engineering technologies have enabled the development of more physiologically-relevant tissue models, which entail a higher degree of complexity, organization and dynamics^26–29^, such as engineered heart tissues (EHTs)^27,30^, isogenic cardiac microtissues^24,31–36^, and organs-on-a-chip^33,34,35–42^.

Organoids, defined as 3D miniaturized versions of an organ, are emerging as promising tools showing realistic micro-anatomy, and organ specific function^9,48–50^. In order to be considered as an “organoid”, an *in vitro* model must fulfil specific requirements, including: (1) 3D multicellular composition with organ-specific cell types, (2) self-organization and histological resemblance to the tissue of origin and (3) recapitulation of at least one specialized biological function similar to the organ being modelled^50–52^.

Well-established organoids have been already generated for brain, kidneys, intestines, guts, lungs and many other organs^9,53^. Until recently, organoid models of the heart had not been established yet^54,55^, as the first such examples only started to emerge in the last couple of years. Notably, early mammalian cardiac organoids showing spontaneous self-organization with distinct atrium- and ventricle-like regions were generated from mouse pluripotent stem cells (PSCs)^56,57^, or as a part of gastruloids^58^. Shortly afterwards, human PSC-derived cardiac organoid models were described^55,59–64^, which were developed with different approaches varying from assembling different cardiac cell types^61^, followed by other self-organized models more faithful to cardiac-specific development^55,62–65^.

Some features of recently reported human cardiac organoids include being modelled after a single chamber of the heart - namely the left ventricle in the case of “cardioids”^55^, relying on an external ECM scaffold, such as Matrigel^60,62^, or featuring the co-emergence of gut tissue together with atrial- and ventricular-like regions^62,64^. While being extremely informative, most of these models lack long-term culture and characterization, which would help them acquire a more mature phenotype, a feature which is usually desirable for more physiologically relevant *in vitro* tissue models^66^.

Here, we aimed to establish induced pluripotent stem cell (iPSC)-based, scaffold-free, long-term human cardiac organoids (hCOs), which self-organize to display discrete atrial and ventricular domains, contain multiple cell types of the human heart, and preserve coordinated contractile activity for several months.

By combining RNA-sequencing and ultrastructural analysis, we demonstrated that dimensionality and time in culture are crucial mediators of survival, differentiation, collective organization and maturation of hCOs.

Finally, we confirmed hCOs qualify as a powerful *in vitro* heart model by proving their response to cardioactive and cardiotoxic drugs in long term culture.

## Results

### Long term 3D culture drives spontaneous self-organization and increased survival in iPSC-derived human cardiac organoids

We aimed to develop a robust differentiation method able to produce cardiac organoids with high reproducibility by using chemically defined culture conditions compatible with drug testing high-throughput analysis.

Since 2D monolayer differentiation of induced pluripotent stem cell (iPSC)-derived cardiomyocytes is very well established in literature, and long-term cultures iPSC-derived cardiomyocyte monolayers naturally tend to delaminate into beating clusters, we first performed human iPSC (hereafter hiPSC) differentiation in a 2D system and test whether they could be used to generate 3D cardiac aggregates in the absence of any external ECM scaffold.

Cardiac differentiation was induced in confluent 2D hiPSC monolayers by sequential modulation of WNT pathway with small molecules, in the absence of insulin^67^ as previously described by Lian et al^13,14^: First, mesoderm specification was achieved by transient WNT activation through chemical inhibition of GSK3, followed by cardiac mesoderm differentiation through the inhibition of the WNT palmitoleoyltransferase PORCN^13,14^ (Figure 1). When beating cell clusters were observed (day 7), insulin supplement was added to the media, since it was needed for cell survival after early contractile cardiomyocytes emerged during differentiation^67^. On day 15 of differentiation, the monolayer was dissociated into single cells and seeded on round-bottom ultra-low attachment plates in order to induce the spontaneous formation of aggregates (Figure 1). Throughout the study, we used 2D monolayer cultures as controls.

**Figure 1.**
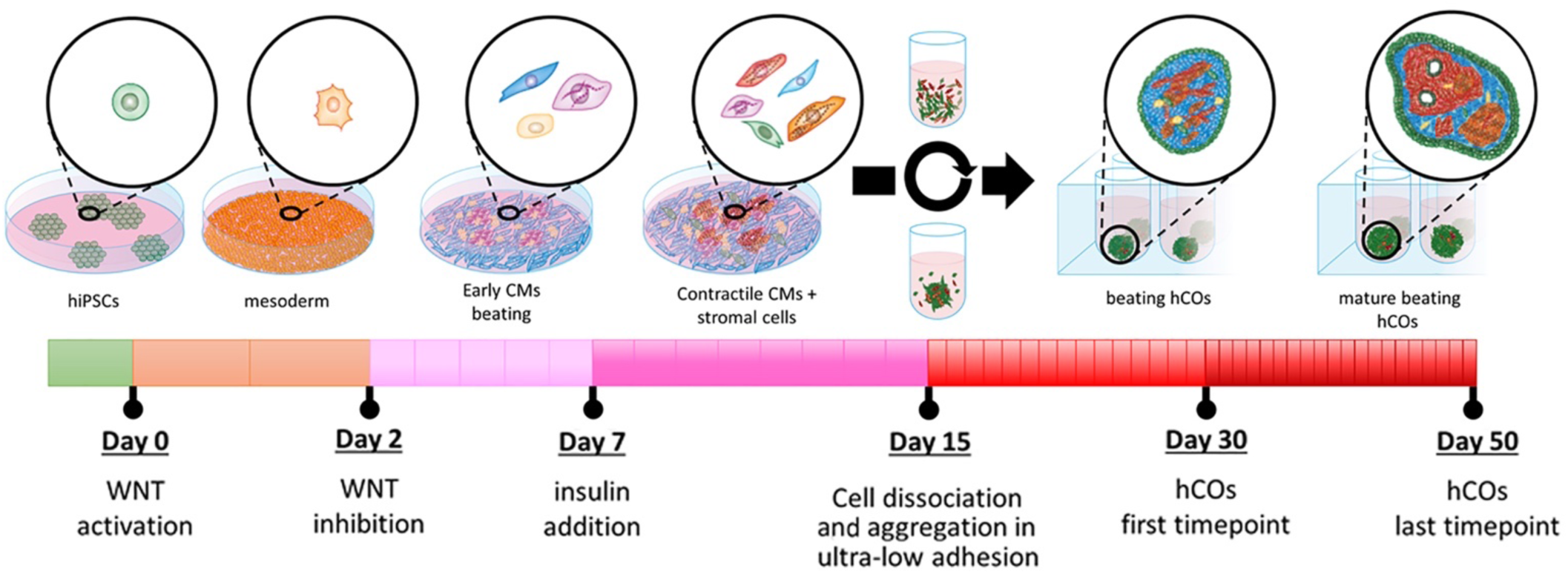
Schematic workflow for the generation and long-term culture of hiPSC derived cardiac organoids. Light green cells represent hiPSCs, red cells represent cardiomyocytes, dark green cells αSMA+ cells, other colours represent other non-myocytes.

After switching the culture from 2D to 3D, we observed the formation of spontaneously beating microtissues. The diameter of the microtissues reached up to 0.9 ± 0.04 mm at day 21 and 1 ± 0.09 mm at day 42 for the long-term culture, in the absence of any external ECM supplementation (Figure 2a, Supplementary video 1 for day 53). The microtissues went on displaying spontaneous contractile activity in long term culture and until at least day 100 (Supplementary video 2 for day 107).

**Figure 2.**
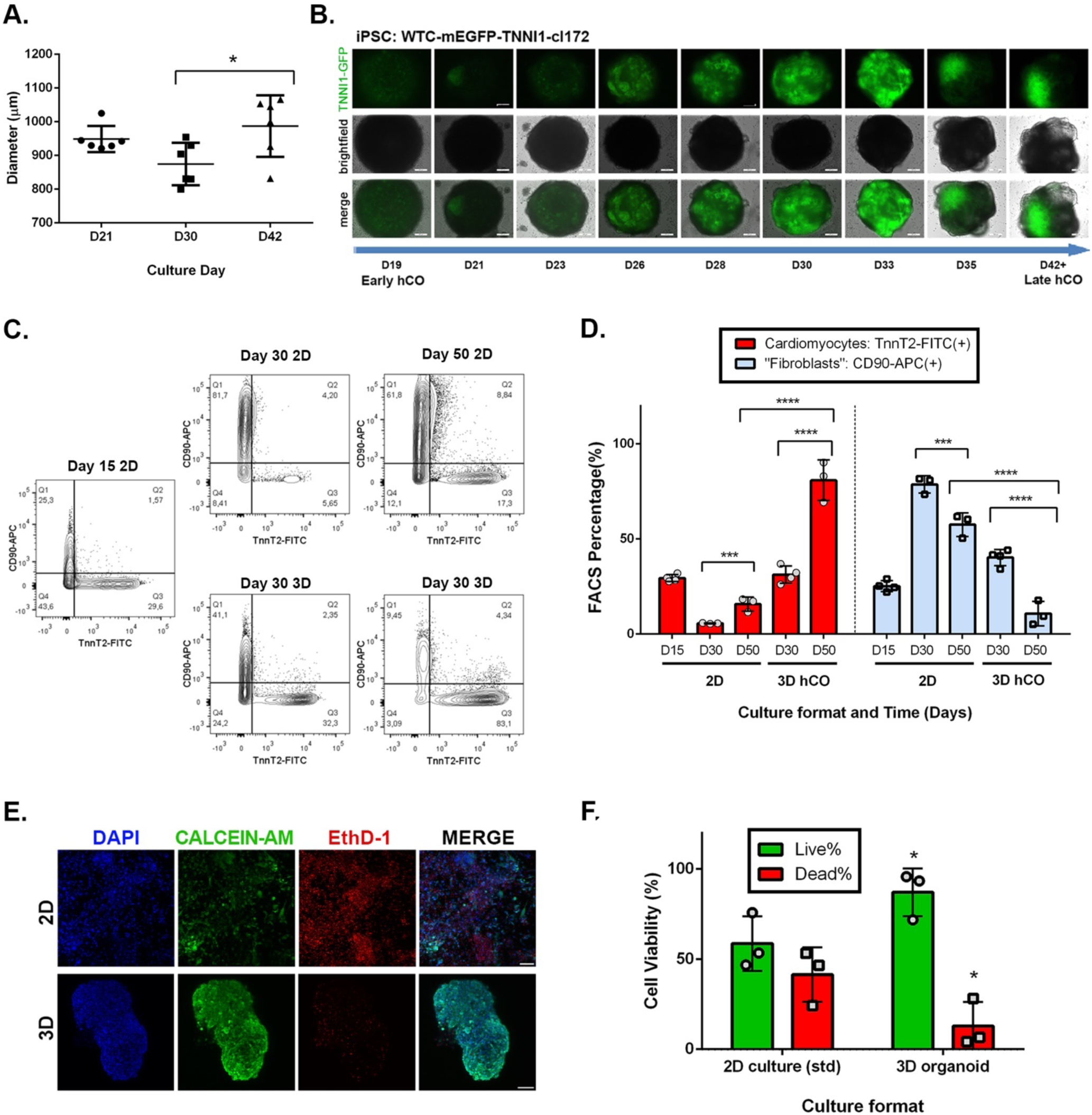
Human Cardiac Organoids Spontaneously Self-Organize and Improve Survival in Long Term Culture. **a**. Average diameter of 3D hCOs over time (n=6, Mean ± SD). **b**. Monitoring spontaneous self-organization of hiPSC-derived cardiomyocytes within 3D organoids expressing GFP-tagged Cardiac Troponin I reporter in hCOs over culture time (Scale bar = 200 μm). **c**. Representative FACS images for of 2D vs 3D on different days of culture, with cardiomyocytes (TNNT2-FITC), and «fibroblasts» (CD90-APC). **e**. Quantification of FACS data with different cell populations with respect to culture dimensionality and time (n=3, Mean ± SD). **e**. Cell viability of 3D hCOs compared to standard 2D cardiac differentiation culture with Calcein/AM staining on day 50, green=live, red= dead. (Scale bar = 200 μm). **f**. Quantification of cell viability on day 50 (n= 3 organoids for 3D, or 3 wells per condition for 2D) (n=3, Mean ± SD)

When the microtissues were kept in culture for longer time (50 days), discrete cellular clusters and chamber-like structures were observed by histological analysis, which were compatible with a phenomenon of self-organization. This feature could not be observed at an earlier timepoint of day 30 (Supplementary figure 1a).

To further monitor this spontaneous cellular organization, we repeated the experiment to generate microtissues from hiPSC expressing GFP-tagged cardiac troponin I (TNNI1) reporter. As expected, we observed the emergence of GFP-expressing contractile cardiomyocytes together with non-tagged cells within the microtissues. With time in culture from day 21 to day 42, the GFP-tagged contractile cardiomyocytes gradually and spontaneously rearranged within the microtissues from a random distribution to one discrete region, and were surrounded by non-GFP tagged cells (Figure 2b).

This observation suggested that our 3D differentiation protocol could give rise not only to cardiomyocytes, but also other cell populations, which showed a tendency to self-organize in extended culture times.

We then quantified the presence of two of the most represented populations in human heart over time ^68,69^ - cardiomyocytes and fibroblasts – in the 3D microtissues by fluorescence-activated cell sorting (FACS) at given time-points (day 30 and day 50 in Figure 2c). The FACS analysis demonstrated that 3D culture is associated with a significant increase in cardiac troponin T2 (TNNT2)-positive cardiomyocytes compared to 2D, especially in long term culture (80.9 ± 10.7 % in 3D day 50 vs 15.7 ± 3.7 % in 2D day 50, p< 0.0001) (Figure 1e). On the contrary, 2D culture was shown to favour the overgrowth of CD90-positive fibroblasts (10.8 ± 6.4 % in 3D day 50 vs 57.5 ± 6.2 % in 2D day 50, p< 0.0001) (Figure 2d and Supplementary figure 1b).

A common drawback of 3D cultures is the possibility that a necrotic area is induced by the poor oxygen and nutrients diffusion towards the core of the construct^70^. In order to rule out the possibility that this was the case in our 3D cultures, we performed a cell viability assay in which we compared 3D hCOs with 2D cultures at the same timepoint (day 50). We found the overall viability of the 3D hCOs to be significantly higher than standard 2D monolayer culture counterpart (87 ± 13.2 % in 3D day 50 vs 58.7 ± 15.1 % in 2D day 50, p< 0.05, Figure 2e and f).

Due to the presence of intrinsic and spontaneous self-organization, cellular heterogeneity, and functional beating in our 3D microtissues over extended culture time, we will hereafter refer to these structures as human cardiac organoids (hCOs).

### Human iPSC-derived cardiac organoids are composed of multiple heart cell types

One of the key features of *bona fide* organoids is the presence of different cell types typical of the given organ, as well as their ability to self-organize in microstructures similar to the organ being modelled^50–52^. In recent reports, the formation of cardiac organoids was associated with the developmental co-emergence of elements of endoderm and mesoderm tissues resembling the development of the heart and gut in the embryo^62,64^.

To explore the spatial distribution of different cell subsets in hCOs, cryosections of the constructs at day 50 of culture were immunostained (IF) to detect the presence of cells expressing markers specific of the different populations in the human heart^69^ (Figure 3). The same slides were also stained for proteins expressed in the gut.

**Figure 3.**
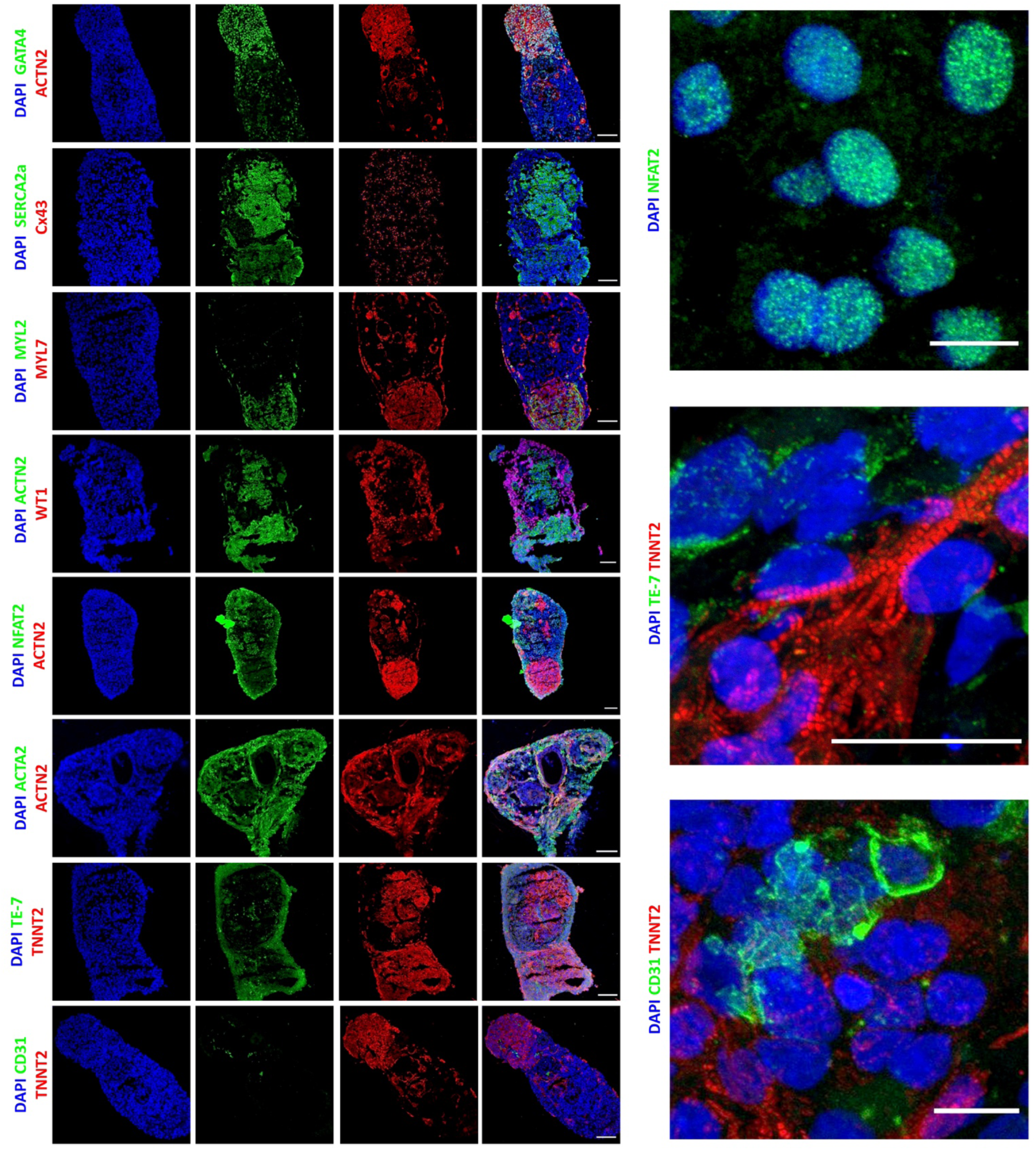
Long term culture of hCOs recapitulates advanced morphology and cellular heterogeneity *in vitro*. Immunofluorescence analysis showing the co-specification of multiple cell type markers in 3D hCO sections on day 50 including cardiomyocytes (ACTN2, TNNT2), atrial (MYL7) and ventricular (MYL2) cardiomyocyte subtypes, epicardial cells (WT1), endocardial cells (NFAT2), smooth muscle/fibroblastic cells (ACTA2), “fibroblasts” (TE-7), endothelial cells (CD31), counterstained with DAPI (blue). Additionally, cardiac morphogenesis (GATA4), calcium ATPase (SERCA2A), and gap junction proteins (Cx43) can be observed. Full organoid scale bar = 100μm, Close-up scale bar = 10 μm.

Confocal imaging showed that cells located at the periphery of the hCOs stained positive for marker of epicardial cells Wilms tumor protein 1 (WT1), fibroblast markers TE-7 and ACTA2 (also known as α-smooth muscle actin, or αSMA), which suggested the presence of a fibrotic shell. The core of the construct was composed by cardiac troponin T2 (TNNT2)- and sarcomeric α-actinin (ACTN2)-positive cardiomyocytes.

Interestingly, we were able to identify two spatially distinct pools of atrial and ventricular cardiomyocytes within the core of the hCOs, by staining with antibodies directed against ventricular (MYL2) or atrial (MYL7) Myosin Light Chain isoforms. Furthermore, markers for cardiac morphogenesis (GATA4), Sarcoplasmic/Endoplasmic Reticulum Calcium ATPase 2 (SERCA2A), and gap junction proteins (Connexin 43, or Cx43) were also observed within the microstructures.

Additionally, the analysis showed the presence of discrete areas which stained positive for the endocardial marker NFAT2, and endothelial cell marker CD31. No markers of embryonic or adult gut tissue, or undifferentiated cell markers were detected in day 50 hCOs (Supplementary figure 2).

Altogether, these results confirmed that hiPSC-derived long term cultured hCOs contain multiple cell types of the human heart organized in functional domains.

### Human iPSC-derived cardiac organoids show ultrastructural organization and maturation in long term culture

A key feature of the adult heart is the existence of a highly recognizable three-dimensional ultrastructure due to the periodical repetition of the functional units of the contractile apparatus, the sarcomere. The length and the alignment of the sarcomeres, together with the interspacing of myosin-actin myofilaments and the abundance and shape of mitochondria, are considered representative of the maturity of the contractile tissue.

We analysed the ultrastructure of the contractile core of the hCOs and assessed how it developed with time in culture by transmission electron microscopy (TEM). TEM analysis clarified that the prototypical contractile apparatus was hardly recognizable in hCOs cultured for 21 days, while highly organized sarcomeres with distinct z-disks and evenly distributed myofilaments could be detected in those cultured for 50 days (Figure 4a, b).

**Figure 4.**
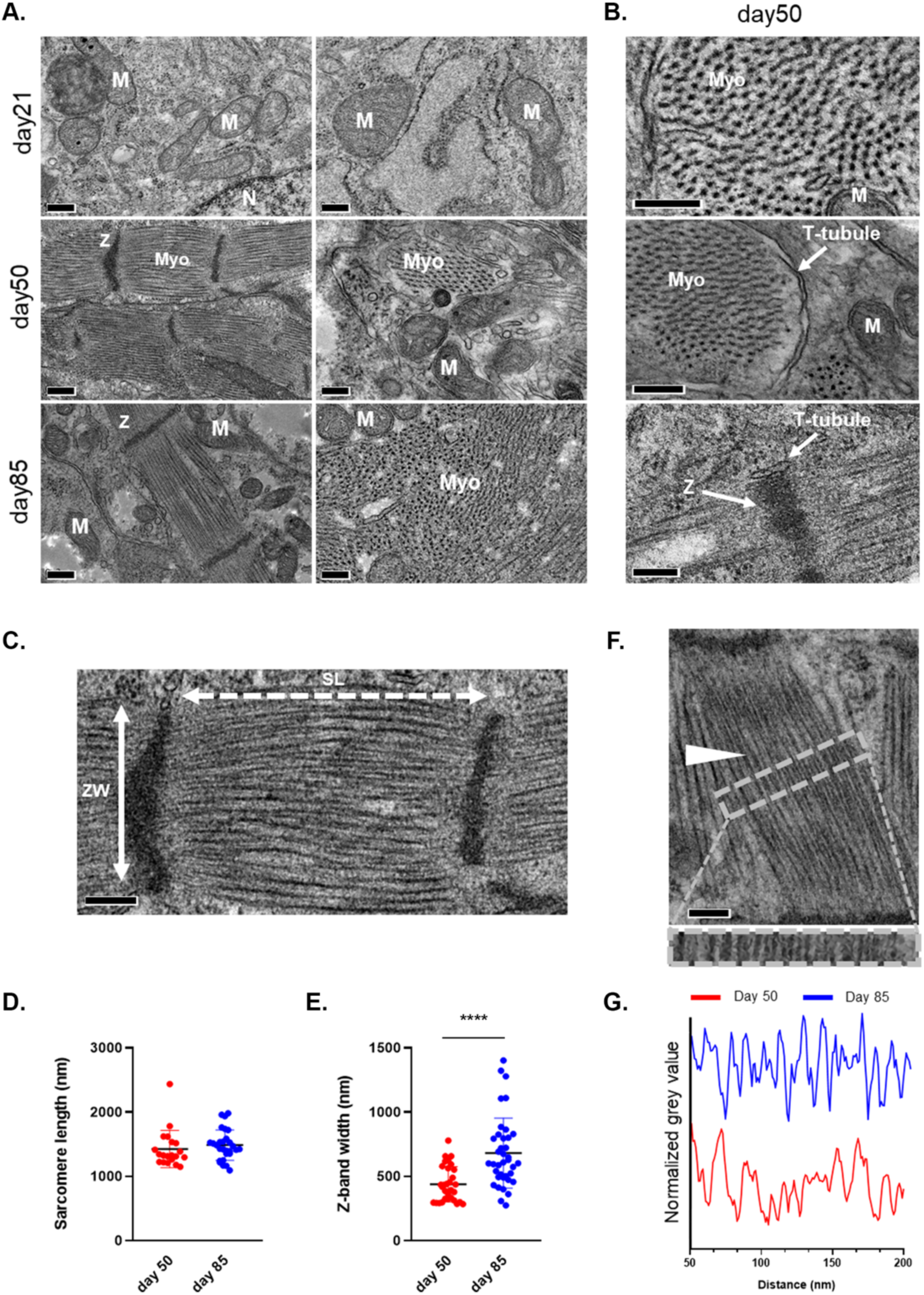
Ultrastructural analysis of hCOs indicates enhanced maturation in 3D environment over time. **a**. TEM images of hCOs showing increasing ultrastructural organization over time, from day 21 to day 85: Cardiomyocyte myofibers (Myo), Mitochondria (M) Z-band (Z), T-tubules. Scale bars = 200 nm. **b**. Close-up images on day 50. Scale bars are 200 nm. **c**. Representative micrograph showing sarcomere length (SL; white dotted two-headed arrow) and z-band width (ZW; cross sectional length; white solid two-headed arrow) with the corresponding sarcomere length. **d**. day 50, 1400 ± 300; day 85, 1500 ± 200; N>20, Mean ± SD)- and z-band width (**e**; day 50, 450 ± 100 nm; day 85 700 ± 250 nm; N>30, Mean ± SD). Scale bar = 200 nm. **f**. Micrograph showing myofiber organization (white arrow) with their relative organization as defined by the plot profile intensity on day 50 vs day 85 of the area indicated by the grey dashed box of “figure 4f”. Scale bar = 200 nm.

The same analysis also demonstrated that longer culture times (day 85) led to the assembly of t-tubules close to the regularly spaced myofibrils, together with the appearance of packed and elongated mitochondria (Figure 4f and Supplementary figure 3). The presence of t-tubules, extensions of the sarcolemma interspersed around the contractile apparatus to maximize the efficiency of calcium exchange, is only found in mature cardiac muscle and was not reported for standard 2D monolayer cultures^71^. In our hCOs, we found the sarcomere length to be approximately 1.5 μm (Figure 4c, d). This value did not change at later time-points (day 85) and is consistent with the values described for young mammalian cardiomyocytes^72^. Meanwhile, Z-band was found to increase slightly in width, but not significantly, with time in culture (700 ± 250 nm on day 85 vs. 450 ± 100 nm on day 50, p <0.0001) as to get closer to the values typical of adult heart^72^ (Figure 4e-g).

Furthermore, TEM also demonstrated a higher amount of mitochondria and glycogen accumulation at later time-points (day 50 and 85) compared to day 21 (Supplementary figure 3), which suggests a metabolic shift to advanced maturation of cardiomyocytes^72^.

All these features indicate hiPSC-derived hCOs undergo structural and metabolic maturation^66^ when cultured for long time in 3D.

### 3D long term culture induces human iPSC-derived cardiac organoid maturation

Contractile cell maturation can be monitored by tracking the evolution of the expression of specific genes encoding for contractile proteins. Hence, we set at investigating the transcriptional landscape of human long-term iPSC-derived 3D hCOs in order to assess the impact of time and dimensionality on the maturation of the contractile cells.

To this end, we performed bulk RNA-sequencing and differential expression analysis (DE analysis) for hCOs at day 30 and 50 of culture and compared them to 2D monolayer cultures harvested at the same time-points.

A total of 2975 genes were found to be significantly and differentially regulated in hCOs compared to monolayer cultures at day 30. This number increased to 6437 at day 50 (Figure 5a and Supplementary tables 1 & 2), possibly indicating a bigger divergence between 2D and 3D cultures over time.

**Figure 5.**
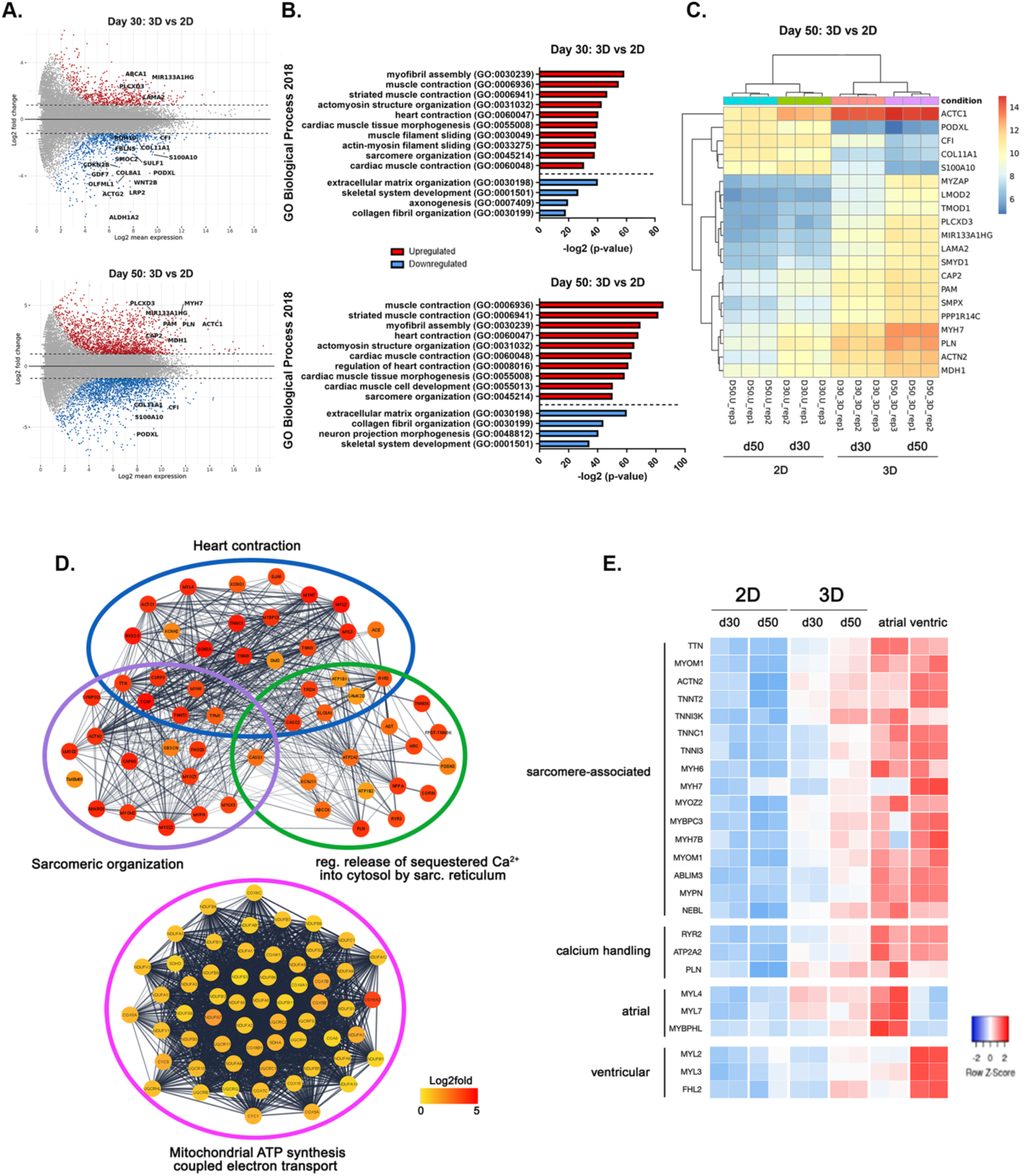
3D long term hCO cultures induce improved cardiac specification and cardiomyocyte maturation at the transcriptional level. **a**. MA plot of the differentially regulated genes between 2D monolayer and 3D hCOs at day 30 (up) and day 50 (down). **b**. Graph representing the -log2value adjusted p-value of significantly upregulated (red) or downregulated (blue) GO Biological process categories when comparing 3D hCOs and 2D monolayers at day 30 (up) and day 50 (down). **c**. Heatmap representing the log2fold change for the top 20 differentially regulated genes between 2D monolayer and 3D hCOs at day 50. **d**. STRING network of the upregulated genes at day 50 hCOs as compared to monolayer cultures for the indicated GO biological processes categories by Cytoscape (Kappa score = 0.3). Log2fold change is color-coded represented for the nodes. **e**. Heatmap encoding for log2fold changes in selected cardiac genes for the 3D hCOs and the 2D monolayers at both time-points in comparison with available datasets for adult human atrial and ventricular tissues.

The functional annotation at both time-points revealed that the genes differentially regulated in 3D hCOs had a fingerprint for cardiogenesis, which included the categories of myofibril assembly and sarcomeric organization, muscle contraction and cardiac tissue morphogenesis (Figure 4b). As an example, the top 20 deregulated genes at day 50 between hCOs and monolayer cultures included genes which are well known to be involved in cardiac muscle maturation and function, namely cardiac muscle α-actin (*ACTC1*), sarcomeric α-actinin (*ACTN2*), phospholamban *(PLN)* and myosin heavy chain 7 (*MYH7*) (Figure 5c). On the contrary, 2D cultures showed an increased expression in ECM-related genes, most likely as a result of the predominance of cardiac fibroblast population in monolayer culture (see Figure 2d and Supplementary figure 4).

Clustering analysis of the genes upregulated in 3D hCOs compared to 2D monolayer cultures at day 50 showed a highly interconnected network of genes involved in heart contraction and sarcomeric organization, which included important structural proteins of the contractile apparatus such as *ACTC1, ACTN2*, troponins (*TNNT2, TNNC1, TNNI1, TNNI3*), myosin heavy (*MYH6, MYH7*) and light (*MYL2, MYL3*) chains, dystrophin (*DMD)*, titin (*TTN)*, obscurin (*OBSCN)* and myozenin 2 (*MYOZ2)*, among many others (Figure 5d).

In good agreement with the presence of t-tubules in hCOs and their drift towards a more mature intracellular Ca^2+^-dependent contraction we detected the enhanced expression of the SERCA Ca-ATPase 2 (*AT2A2*) and its inhibitor phospholamban (*PLN*), triadin (*TRDN*), ryanodine receptors (*RYR2* and the Purkinje’s cell specific *RYR3*), calsequestrins (*CASQ1, CASQ2*), as well as genes related to cardio-renal homeostasis angiotensinogen, atrial natriuretic peptide and corin (*AGT, NPPA, CORIN*). Interestingly, in accordance with the marked increase in the number of mitochondria observed in the TEM images (see Figure 4a), day 50 hCOs showed an upregulation in the network of genes associated to ATP synthesis coupled electron transport, including several NADH:ubiquinone oxidoreductase and Cytochrome c oxidase subunits.

Since long term 3D cultures seemed to be able to promote the maturation of hCOs, we set at comparing the transcriptomic landscape of our *in vitro* constructs with available datasets obtained from adult atrial and ventricular heart tissues^73,74^ (Figure 2d, Supplementary figure 4). We figured using these datasets might help us confirm the results of our confocal analysis hinting at the presence of distinct atrial and ventricular populations in our hCOs (see Figure 3).

Sample clustering showed that day 50 hCOs were closer to adult heart at transcriptional level, with the most similar levels being associated to the expression of several sarcomere-associated genes, including *TTN, ACTN2*, troponins (*TNNT2, TNNI3K, TTNC1, TNNI3*) and Myosin heavy chains (*MYH6, MYH7, MYH7B*), as well as important components of the calcium handling apparatus (*RYR2, ATP2A2, PLN*). Consistent with our immunostaining results, day 50 hCOs displayed both atrial (*MYL4, MYL7, MYBHL*) and ventricular (*MYL2, MYL3, FHL2*) chamber markers (Figure 5e).

Overall, our transcriptomics analysis confirmed that, in our 3D hCOs, culture dimensionality is a crucial factor mediating hiPSC-derived cardiac specification and cardiomyocyte maturation at a metabolic, calcium handling and sarcomeric level. The maturation of 3D hCOs is strongly promoted by time in culture.

### Human iPSC-derived cardiac organoids functionally respond to drugs in a dose dependent fashion in long term culture

An essential feature of *bona fide* organoids is to replicate at least one specialized function of the modelled organ^50–52^, which in the case of heart tissue can be assessed by the contractile activity of hCOs. As previously indicated, once the hCOs acquired spontaneous contractile activity early in culture, the contractility persisted for more than 100 days (Supplementary video 2).

In order to test their physiological significance, hCOs were exposed to clinically relevant doses of cardioactive drugs^75,76^ in long-term cultures (> day 50). In detail, isoproterenol and verapamil were used as positive and negative inotropes, respectively. Isoproterenol, a beta-adrenergic agonist, increases the contractile force and beating frequency^77^, while verapamil, a calcium channel blocker, decreases the beating rate^78^. The hCOs were incubated with increasing drug doses (0.01 - 1μM) for 15-20 minutes and further monitored by live imaging using a confocal microscope.

The response to the treatment was evaluated and quantified based on the contraction (Figure 6a) and beating rate (Figure 6b) of the hCOs, analysed through the open-source software tool MUSCLEMOTION^79,80^. The acquired data shows that increasing concentrations of isoproterenol resulted in enhanced contraction amplitude and beating rate, with the beating rate peak at 1 μM. As expected, increasing concentrations of verapamil had the opposite effect, with the beating being completely stopped at 1 μM drug dosage (Supplementary videos 3-8).These observations are in accordance with those reported in previous studies in cardiac microtissues^81^ and confirm that hCOs can functionally respond to cardioactive drugs in a dose dependent manner.

**Figure 6.**
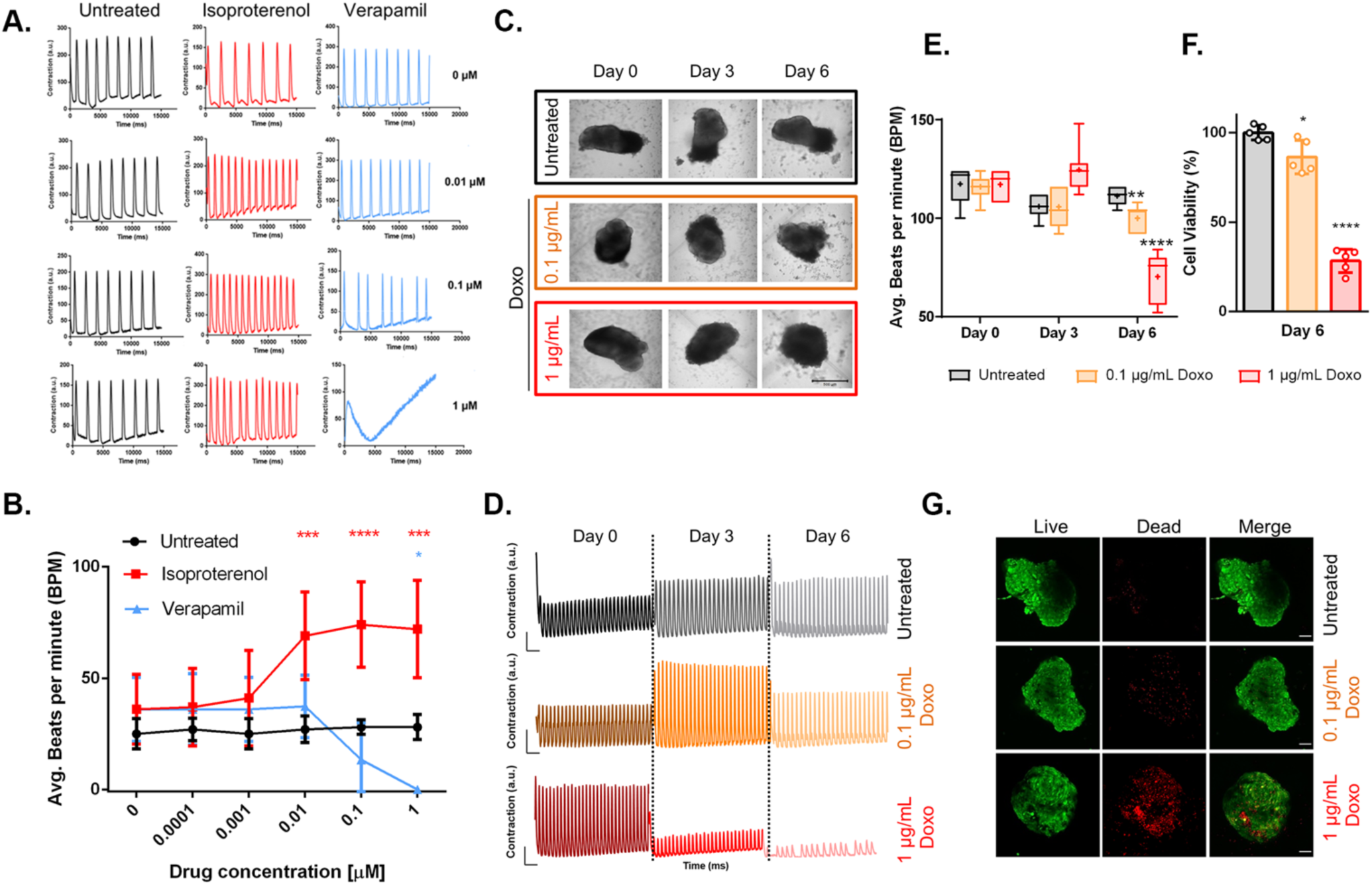
3D hCOs show functional response to cardioactive and cardiotoxic drugs in a dose and time dependent manner. **a**. Representative contraction amplitude plots of long-term (>day 50) hCOs in response to untreated vs increasing doses of Isoproterenol and Verapamil. **b**. Average beating rates of hCOs in response to untreated vs increasing doses of Isoproterenol and Verapamil. (n=3, Mean ±SD) **c**. Morphology of long-term (>day 50) hCOs in response to untreated vs increasing doses of Doxorubicin (Doxo) over 6 days. (Scale bar = 500 μm). **d**. Representative contraction amplitude plots of hCOs in response to untreated vs increasing doses of Doxo over 6 days (t=15s). **e**. Average beating rates of hCOs in response untreated vs increasing doses of Doxo over 6 days. (n= 6, Mean ±SD) **f**. Luminescence-based quantification of cell viability of 3D hCOs on day 6 of Doxo treatment normalized to untreated samples (untreated vs 0.1 μg/mL vs 1μg/mL) (n= 6, Mean ±SD). **g**. Representative cell viability of hCOs on day 6 of Doxo treatment (untreated vs 0.1 μg/mL vs 1μg/mL) with Calcein/AM staining, green =live, red= dead. (Scale bar = 100 μm)

Finally, we investigated the possibility of modelling drug-induced cardiotoxicity on hCOs, by treating them with doxorubicin (Doxo), a well-known chemotherapeutic agent widely used in the treatment of several types of cancers^82^. The drug is - in fact - also known to have cardiotoxic and pro-fibrotic effects, thus able to cause or exacerbate heart failure *in vivo*^83–88^.

hCOs were exposed to 0.1 μg/mL and 1 μg/mL of Doxo and their morphology and contractile activity was compared with untreated controls (n=6) for the next following 6 days. The culture medium was exchanged every 3 days and hCOs monitored on a daily basis. Figure 6c shows that the morphology of hCOs gradually changed over the course of treatment, and with the organoids displaying more irregular edges when exposed to higher Doxo doses. In addition, clear changes could also be observed in the beating profiles of hCOs treated with Doxo (Figure 6d, e, and Supplementary videos 9-14).

We figured the changes in the morphology of hCOs treated with the chemotherapeutics might be due to the induction of cell death by the drug. Therefore, after 6 days of Doxo treatment, we assessed the hCOs viability with a luminescence-based assay. The results in Figure 6e, f revealed that the hCOs treated with 1 μg/mL Doxo were significantly less viable than both the control group and the group treated with lower drug concentration (0.1 μg/mL) (Average viability on day 6: 100 ± 4.1% for ctrl, 86.4 ± 9.3 for 0.1 μg/mL Doxo, 28.3 ± 6,6% for 1 μg/mL Doxo, p<0.0001)

## Discussion

In this study we present a straightforward method to generate complex and functional 3D iPSC-derived human cardiac organoids in the absence of an external scaffold.

Our study has demonstrated that long-term cultured 3D hCOs have structural and functional characteristics resembling native human heart such as: (1) spontaneous self-organization; (2) presence of multiple cell types of the heart belonging to epicardium, myocardium and endocardium layers; (3) ultrastructural organization and maturation of cardiac muscle sarcomeres and mitochondria; (4) distinct atrial and ventricular regions; (5) improved survival, cardiac specificity and maturation at transcriptional level; (6) long-term functional beating in the absence of external stimuli; (7) functional response to cardioactive and cardiotoxic drugs in a dose- and time-dependent fashion.

When compared to standard 2D cardiomyocytes culture, 3D hCOs showed an enhanced survival rate. More importantly, their cellular composition appeared to be more representative of the physiological conditions in the heart, although their overall phenotype appeared to be close to foetal heart^81,89^.

While the effects of long term culture in promoting the maturation of PSC-derived cardiomyocytes is well established^90,91^, 2D culture is usually challenging in the long run, as the cardiomyocytes tend to delaminate from the traditional tissue culture plates^90^, leaving non-myocyte cell populations such as fibroblasts to overgrow.

One of the greatest advantages of our model is that there is no need for an external scaffold as the hCOs self-organize and survive for a long time in static culture conditions. Other models have been developed in the absence of an external ECM, however they usually require the differentiation of other stromal cells (such as epicardial or fibroblastic cells) separately, followed by their assembly with cardiac myocytes ^34,55^.

On the contrary, by following the original 2D cardiac differentiation protocol by Lian et al^13,14^ and simply switching the dimensionality from 2D to 3D, we have generated long-term 3D hCOs in which epicardial cells, fibroblasts, and endocardial cells emerge spontaneously together with cardiomyocytes, without the need to set up separate lineage differentiation protocols and further assembly. In addition, the spontaneous beating of our hCOs was observed for more than 100 days as, to this date, only reported by Silva et al. as well. The latter paper, though, reported that the developmental co-emergence of gut also occurred together with the heart tissue^64^.

In our model, a distinct gut tissue was not observed after 50 days of culture through immunofluorescence analysis. Histological analysis of the hCOs has shown distinct populations of WT1-positive epicardial cells, TE-7-positive fibroblast-like cells, NFAT2-positive endocardial cells along with a distinct core of cardiomyocytes. CD31-positive endothelial cells could also be detected, however at almost negligible levels. This finding might be explained by the fact that in native human heart endothelial cells have different developmental origins^92^, and we did not supplement our culture media with endothelial-lineage favouring factors, like VEGF, during our study.

Another remarkable feature of the hCOs is the representation of distinct atrial and ventricular cardiomyocyte populations, which were clearly identified by confocal imaging of MYL2 and MYL7. Although with significant differences, a similar phenomenon of atrial/ventricular cardiomyocyte subtype distinction was also shown by Israeli and colleagues, which appeared to be blunted with long term culture^63^. Instead, the herein reported hCOs revealed higher cardiac-specific cellular heterogeneity and maturation in terms of histological, ultrastructural, and transcriptional-levels in long term 3D cultures compared to shorter culture times and standard 2D monolayer culture.

The superior maturity and complexity of our long-term 3D hCOs could be also demonstrated at the ultrastructural level by our TEM analysis over long culture times. At day 50 and 85, our hCOs displayed clearly aligned myofibers resulting in well-organized sarcomeres, as compared to earlier time points. These morphological adaptations, which are hallmarks of mature cardiac tissues^72^, were further supported by transcriptomics analysis, where hCOs showed an increased expression of genes related to cardiac contraction and sarcomeric structures.

The electrophysiology and calcium handling properties of immature cardiomyocytes diverge in crucial ways from their mature counterpart. While immature cardiomyocytes can rely on the calcium released at the cell periphery to initiate sarcomere contraction, mature cardiomyocytes develop plasma membrane invaginations (t-tubules) which are wrapped around the myofibrils as to make the process of calcium release and re-uptake faster and more efficient^93,94^. The first polarization triggers the release of calcium stored in the sarcoplasmic reticulum by RYR2, which is back to the sarcoplasmic reticulum by SERCA2 (sarco/endoplasmic reticulum Ca^2+^-ATPase). The analysis of hCOs confirmed the presence of t-tubules at the longer time-points (> day 50), the upregulation of some of the genes involved in their biogenesis (*JPH2, ACTN2, NEXN*) and sarcomeric calcium management (*RYR2, PLN, SERCA*), hence demonstrating long-term 3D culture determines functional maturation of hCOs.

Finally, cardiomyocytes also experience several metabolic adaptations during *in vivo* maturation in order to generate the required amount of ATP^95^. Our ultrastructural results are compatible with this metabolic switch, showing that hCOs acquired elongated mitochondria distributed around the contractile apparatus with time in culture. This idea is further reinforced by the upregulation in long term 3D cultured hCOs of genes related to chain electron transport-linked phosphorylation.

Functional analysis of the 3D hCOs proved that they were responsive to clinically relevant doses of cardioactive and cardiotoxic drugs. This is the first step required to consider the generation of patient-specific models of disease for drug testing.

As for modelling chemotherapy-induced cardiotoxicity, while doxorubicin is a well-established chemotherapeutic known to cause heart failure as a side effect^83–88^, the *in vitro* modelling in cardiac microtissues is still not well-established due to the limitations so far encountered in reproducing the physiological cellular heterogeneity, and cardiomyocyte maturity^61^. Furthermore, it is crucial to consider the influence of cell-cell interactions of cardiomyocyte vs non-myocytes - especially fibroblasts - and cell-ECM interactions^96^. Our hCOs platform offers the advantages of long-term viability, cellular heterogeneity and advanced maturity *in vitro*, rendering it a physiologically relevant alternative for future studies. For instance, further clinical parameters such as the cardio-fibrotic potential of doxorubicin treatment^86,87,97^, the secretion of cardiac troponins and natriuretic peptides as clinically relevant markers of chemotherapy-induced cardiotoxicity^98^ or simply the potential for personalized medicine could be exploited by using this iPSC-derived hCOs^99^.

Nevertheless, despite the remarkable structural and functional maturation in the long-term culture model as compared to traditional monolayers, our hCOs still displays features of the foetal rather than the adult heart. For example, myocytes of our hCOs have ∼1.5 μm length of sarcomeres (as in foetal mammalian hearts), whereas in adult hearts, this length is usually 2 μm^72^. We believe the adoption of additional stimuli (i.e.: dynamic culture, mechanical conditioning and electrical stimulation) might in the future help further improve the model here described^100^.

In a longer perspective, *in vitro* cardiotoxicity studies should not underestimate the importance of multi-organ interactions. While drugs in their native chemistries may not always result in direct cardiac damage^101^, their metabolic by-products might cause cardiotoxicity *in vivo*^101,102^.

In conclusion, we have established a simple procedure to prepare human induced pluripotent stem cell-derived cardiac organoids recapitulating some key features of the human heart, and displaying improved maturation and functionality in long term culture, thus enhancing the potential of *in vitro* platforms to be used in translational research applications.

## Methods

### Stem cell culture and maintenance

The human iPSC cell line DF 19–9-7 T (iPS, karyotype: 46, XY) was purchased from WiCell (Madison, WI, USA), and the Troponin I1 reporter iPSC line (TNNI1-iPS) was purchased from Coriell Institute (Cat.no. AICS-0037-172, Camden, New Jersey, USA). All iPSC cells were cultured and maintained in feeder-free conditions as previously described^103,104^ on Growth Factor Reduced Matrigel®-coated plates, (1:100 in DMEM/F12, Corning) in complete Essential 8™ Medium (E8, Thermo Fisher Scientific) containing penicillin/streptomycin (P/S) (0.5%, VWR), incubated at 37°C, 5% CO_2_.

### hiPSC-derived monolayer cardiac differentiation and maintenance

Cardiac differentiation from hiPSCs was performed following the Wnt signalling modulation protocol by Lian et.al.^13,14^, with slight modifications as previously described^103^. Briefly, prior to cardiovascular differentiation, hiPSCs were dissociated into single cells (TrypLE Select, Thermo Fisher Scientific) and re-seeded onto 12-well Matrigel-coated plates with 2.0 × 10^5^ cells/cm^2^, with complete Essential 8 medium including Y27632 Rock Inhibitor (RI) (1:4000 dilution from 10 μM stock, 2.5 μM final, Selleck chemicals, Houston, TX, USA). The next day, the medium was replaced with complete Essential 8 medium without RI, and the medium exchange was performed daily until the cells reached 100% confluency. On day 0, to start mesoderm differentiation, the medium was changed with RPMI 1640 with L-Glutamine (Biosera) media supplemented with P/S, B-27™ supplement minus insulin (1×, Thermo Fisher Scientific) and CHIR99021 (8 μM, Sigma-Aldrich). On day 2, the medium was exchanged with RPMI 1640+B-27 minus insulin (RPMI+B27-Ins), supplemented with IWP-2 (5 μM, Selleck chemicals). On day 4, the medium was replaced with RPMI+B27-Ins, and medium exchange was performed every 2 days until the cells started beating (usually on day 7, but may depend on the iPSC line). Once beating clusters started to emerge, the medium was replaced with RPMI 1640+B-27™ supplement (RPMI+B27+Ins) (1×, Thermo Fisher Scientific), and medium exchange was performed every 2-3 days until day 15 of differentiation. From day 15 on, the cells were used either for generation of 3D hCOs, or were continued to be cultured as 2D monolayers as controls, with medium exchange every 3-4 days until the end of experiments.

### hiPSC-derived cardiac organoid generation and maintenance

On day 15 of monolayer cardiac differentiation, cells were dissociated by putting 0.5 mL StemPro™ Accutase™ (Gibco) per well, and incubating in room temperature or in 37°C incubator for 10-20 minutes with occasional mechanical pipetting, until the cells were visually dissociated. After sufficient dissociation, Accutase™ was stopped by putting 3mL RPMI 1640 with 20% Knockout Serum Replacement (Gibco) per well, and the cells were pelleted by centrifugation at 150xg for 3 minutes. The resulting pellet was resuspended with RPMI/B27+Ins + P/S + RI (1:2000 dilution from 10 μM stock, 5 μM final), counted by LUNA™ cell counter (Logos), and seeded 150,000 cells/well in 96-well round bottom ultra-low attachment plates (Corning Costar 7007) at a volume of 150 μL/well. The plates were then centrifuged at 200xg for 5 minutes and incubated at 37°C, 5% CO_2_. As 2D control, either 200,000 cells were seeded in Matrigel-coated 96-well tissue culture plates at a volume of 200 μL/well, or 1.5 × 10^6^ cells were seeded on Matrigel-coated 24-well tissue culture plates in 1 mL volume per well, or left undetached in 12-well plates. After 48 hours, medium was replaced with RPMI/B27+Ins without RI from hCOs and 2D-controls that were re-seeded. From this point on, medium was changed partially every 2-3 days with RPMI/B27+ Ins +P/S until day 30, then every 3-4 days at least until day 50, or longer.

### Cell viability with Calcein AM/EthD-1 staining

3D hCO and 2D culture control viability was assessed with LIVE/DEAD™ Viability/Cytotoxicity Kit, for mammalian cells (Thermo Fisher Scientific), according to manufacturer’s instructions, and confocal imaging was performed with Zeiss LSM 780 confocal microscope, the viability was later quantified by image processing with ImageJ (v1.53o).

### Immunofluorescence (IF)

Organoid fixation, cryosectioning, and immunostaining was modified from Perestrelo et al^96^. Briefly, 3D hCOs were collected with low binding pipette tips, and fixed in 4% PFA (Santa Cruz Biotechnology) supplemented with 0.03% eosin (Sigma Aldrich) for 2 hours at room temperature, followed by washing with 1xPBS (Lonza), and embedding in 30% sucrose solution at 4C until they sink at the bottom (∼2 days). The organoids were then embedded in OCT solution (Leica), frozen in cassettes embedded in isopentane (VWR) cooled with dry ice, and were stored in -80°C until further processing. The frozen organoids were cryosectioned by cryotome (Leica CM1950) onto Menzel Gläser, SuperFrost® Plus slides (Thermo Fisher Scientific) at 10μM thickness.

For IF analysis, the organoid sections were washed with PBS for 2×5 min at RT, followed by permeablization with 0.2% Triton X-100 (Sigma Aldrich) for 5 min. Blocking was done by 2.5% bovine serum albumin (BSA) (Biowest) in PBS for our at room temperature. Primary antibodies were incubated overnight at 4°C (Supplementary table 3). The next day, the samples were washed with PBS, followed by appropriate Alexa-conjugated secondary antibodies (Supplementary table 3). Counterstaining was done by DAPI (Roche), and organoid slides were mounted with Mowiol®4-88 (Sigma Alrdich). Confocal imaging was performed with Zeiss LSM 780 confocal microscope.

Further histological procedures can be found in Supplementary methods.

### Organoid dissociation

For FACS and RNA isolation, organoids were collected in Eppendorf tubes with low binding pipettes on day 30 and 50, and dissociated with MACS MultiTissue Dissociation Kit 3 (Miltenyi Biotec) according to manufacturer’s instructions. As controls, 2D monolayer cultures were directly dissociated on the plates with the same kit.

### Microscopy and image analysis

Confocal imaging was performed with Zeiss LSM 780 confocal microscope. Light microscopy imaging was performed with Leica DM IL LED inverted microscope, and fluorescence images were collected with Leica DMI4000 B inverted fluorescence microscope. Image analysis was performed with ImageJ (v1.53o).

### RNA isolation, bulk RNA-sequencing and DE analysis

For RNA isolation, 3D hCOs and 2D controls were dissociated by using MACS MultiTissue Dissociation Kit 3 on day 30 and day 50. After cell dissociation, RNA was extracted with High Pure RNA Isolation Kit (Roche) according to manufacturer’s instructions, and quantified with Nanodrop 2000 Spectrophotometer (Thermo Fisher Scientific). Quality control for RNA integrity number (RIN) was also determined with Agilent 2100 Bioanalyzer.

Bulk RNA-sequencing and DE analysis procedure is further described in Supplementary methods.

Gene ontology (GO) enrichment analysis for biological processes was performed with EnrichR web-tool^105–107^. Clustering analysis of upregulated genes was done with Cytoscape^108^.

RNA-seq data from this study were also compared to adult human heart RNA-seq data from BioProjects PRJNA667310^73^ and PRJNA628736^74^ obtained from the NCBI BioProject database (https://www.ncbi.nlm.nih.gov/bioproject/) (Accession numbers: SRR12771050, SRR12771052, SRR11620685, SRR11620686) by using Biojupies web-tool^109^.

### Fluorescent activated cell sorting (FACS)

Cells dissociated with Accutase on Day 15 of differentiation, and with MACS Multi-tissue dissociation kit 3 (Miltenyi Biotec Ca No. 130–110-204) at day 30 and 50, along with corresponding 2D controls were analysed by Beckman Coulter MoFlow Astrios Cell Sorter (Beckman Coulter Life Sciences) as described previously ^96,104^, with CD90-APC, TNNT2-FITC antibodies, and their corresponding unstained controls(Supplementary table 3). Cell populations were quantified by using FlowJo software V10 (Tree Star).

### Transmission Electron Microscopy (TEM) sample preparation and analysis

hCOs were transferred to new well-plates with low binding pipettes, and washed 2 x with PBS. Fixation was done by 1.5% PFA+ 1.5% glutaraldehyde (Sigma Aldrich) diluted in RPMI 1640 culture media for 1 hour, then washed with 0.1 M cacodylate buffer (pH 7.4) (Sigma Aldrich) for 3 times 5 minutes each, and incubated overnight at 4°C in cacodylate with glutaraldehyde. The next day, samples were rewashed 3 times with cacodylate buffer, and incubated at 4°C in cacodylate buffer until sending the samples, protected from light. The organoids were post-fixed for 1.5 hours with 1% osmium tetroxide in 0.1 M cacodylate buffer and washed for 3 times with 0.1 M cacodylate buffer, 10 minutes each, then stained with 1% uranyl acetate in milli-Q water overnight at 4°C, followed by washing in milli-Q water. The samples were then dehydrated in an ascending EtOH series using solutions of 70%, 90%, 96% and 3 times 100% for 10 minutes each, incubated in propylene oxide (PO) 3 times for 20 minutes before incubation in a mixture of PO and Epon resin overnight, and incubated in pure Epon for 2 hours and embedded by polymerizing Epon at 68°C for 48 hours. Ultra-thin sections of 70 nm were cut using a Leica Ultracut EM UC 6 Cryo-ultramicrotome. TEM images were collected with a JEOL JEM 1011 electron microscope and recorded with a 2 Mp charge coupled device camera (Gatan Orius).

### Drug treatments

Isoproterenol hydrochloride (Sigma Alrdich) was dissolved in dimethyl sulfoxide (DMSO) (Sigma Alrdich), and Verapamil hydrochloride (Sigma Alrdich) in methanol (VWR) to prepare 1 M stock solutions. Each drug solution was 10-fold serially diluted in RPMI/B27+Ins media to make final concentrations between 0.0001-1 μM. As untreated controls, complete RPMI/B27+Ins media supplemented with DMSO or methanol were used. For data collection, the hCOs with drug-supplemented media were incubated for 10 minutes at 37°C, 5% CO_2_ each time the drug concentration changed, followed by live imaging with video acquisition per condition with Zeiss LSM 780 confocal microscope. Chemotherapy-induced cardiotoxicity was evaluated in hCOs treated with RPMI/B27+Ins supplemented with doxorubicin (Doxo) (Sigma Aldrich), at 0.1 μg/mL and 1 μg/mL concentrations, or plain RPMI/B27+Ins as control for 6 days. Growth medium was exchanged every 3 days, and organoid morphology and beating were monitored daily by live video recording with Zeiss LSM 780 confocal microscope for 6 days. On day 6 of drug treatment, the organoids were collected for evaluating viability.

### Live video acquisition and contraction analysis

To evaluate the contractile function of organoids, live image sequences were acquired with Zeiss LSM 780 Confocal microscope using transmitted light mode (37°C, 5% CO_2_), as equivalent to 60 fps for 15 seconds; where the image sequences were later turned into .avi files with ImageJ. The beating rates were counted and calculated manually from the video analysis. and representative contraction graphs were derived with the open-source video analysis software MUSCLEMOTION ^79,80^ per providers’ instructions.

### Luminescence-based cell viability assay

hCO viability after 6 days of doxo treatment was assessed by CellTiter-Glo® Luminescent Cell Viability Assay Kit (Promega) according to manufacturer’s instructions. After the hCOs were dissociated by the assay treatment, the samples were transferred to a clear-bottom 96-well white assay plate (Corning Costar 3610) and cell viability was quantified with Centro LB 960 Microplate Luminometer (Berthold Technologies).

### Statistics

Data are expressed as mean ± SD. Statistical analysis was performed using Microsoft Excel 2010 and GraphPad Prism v6.01 using one-way ANOVA (with Sidak’s multiple comparisons tests or Tukey’s multiple comparisons test) and two-way ANOVA tests (with Sidak’s multiple comparisons test or Dunnett’s multiple comparisons test), where p<0.05 was considered to be significantly different as denoted with asterisks [(*) p ≤ 0.05, (**) p ≤ 0.01, (***) p ≤ 0.001, (****) p ≤ 0.0001]. Sample sizes of independent experiments were described in the figure legends where applicable.

## Data availability

The RNA-sequencing data obtained and presented in this study were generated at CF Genomics of CEITEC, and the dataset will be freely available after getting an accession number. The data that support the findings of this study are available upon reasonable request.

## Acknowledgements

We thank Václav Hejret, Jana Vašíčková, and Jana Bartoňová for scientific advice and technical assistance. We acknowledge the CF Genomics of CEITEC supported by the NCMG research infrastructure (LM2018132 funded by MEYS CR) Bioinformatics for their support with obtaining scientific data presented in this paper. We are grateful to Ivana Andrejčinová., Marco de Zuani and Jan Frič for the NFAT2 and ASCL1 antibodies. We also thank Romana Vlčková, Hana Duľová, and Helena Ďuríková for their support on continuation of the study.

Giancarlo Forte, Stefania Pagliari and Vladimír Vinarský were supported by the European Regional Development Fund - Project ENOCH (No. CZ.02.1.01/0.0/0.0/16_019/0000868). Jorge Oliver De La Cruz and Soraia Fernandes were supported by the European Social Fund and European Regional Development Fund-Project MAGNET (CZ.02.1.01/0.0/0.0/15_003/0000492). Marco Cassani, an iCARE-2 fellow, has received funding from Fondazione per la Ricerca sul Cancro (AIRC) and the European Union’s Horizon 2020 research and innovation programme under the Marie Skłodowska-Curie Grant Agreement No. 800924. Francesco Niro and Daniel Sousa have received funding from the European Union’s Horizon 2020 research and innovation programme under grant agreement No 860715.

## Author contributions statement

E.E. conceptualized the study, designed and performed the experiments, analysed the data and drafted the manuscript. J.O.D.L.C., S.F., M.C., V.V., A.R.P. and S.P. provided essential information and techniques, and assisted in designing, planning, carrying out and interpreting experiments. F.N. and D.S. performed experiments and analysed the data. J.V. and J.O.D.L.C. assisted with bioinformatics analysis. M.C. and D.D. performed the TEM analysis. F.C. and H.R. assisted in funding acquisition, and manuscript revision. P.E. provided study supervision and manuscript revision. G.F. provided study supervision, funding acquisition, paper conceptualization and manuscript finalization. All authors contributed in interpretation, and presentation of data as well as manuscript editing. All authors reviewed and approved the final version.

## Competing interests

None

## List of supplementary materials

Supplementary video 1– Representative beating hCO

Supplementary video 2 – Day 100+ organoid video

Supplementary Figures:

*SI1– a. H&E staining and IF on D30 vs D50 hCO, b. 2d monolayer vs 3d hCO on day 50*
*SI2 – IF hCOs for undifferentiated cells & gut tissue*
*SI3 - Supplementary figure 3 – Additional TEM images*
*SI4 – Supplementary transcriptomics analyses for differentially regulated genes between day 30 and day 50 of culture for 3D hCOs and 2D monolayer culture*.

Supplementary Methods – Histology (H&E), RNAseq and DE analysis.

Supplementary videos (3-8) – Beating hCOs before & after max dose of cardioactive drugs

Supplementary videos (9-14) – Beating hCOs before & after doxorubicin on Day 0 and Day 6

Supplementary table 1 – log2changes and p-values for the different comparisons

Supplementary table 2 – GO BP clustering analysis for the differentially regulated genes at the different comparisons

Supplementary table 3 – List of antibodies used in the study (IF + FACS)

## Notes

### Competing Interest Statement

The authors have declared no competing interest.

