## Supplementary table 3 for "Generation and Maturation of Human iPSC-derived Cardiac Organoids in Long Term Culture"

| **Immunofluorescence** | | | | |
| --- | --- | --- | --- | --- |
| ***IF- Primary Antibodies*** | | | | |
| **Name** | **Host species** | **Dilution** | **Company** | **Catalog no** |
| Anti-WT1 | Rabbit | 1:200 | Cell Signalling | 83535 |
| Anti-ACTA2 (α-SMA) | Rabbit | 1:500 | Abcam | ab5694 |
| Anti-ACTN2 (sarcomeric α-actinin) | Mouse | 1:800 | Sigma Aldrich | A78811 |
| Anti-ASCL2 | Sheep | 1:50 | Thermo Fisher Scientific | PA5-47852 |
| Anti-Brachyury (T) | Goat | 1:50 | R&D Systems | 967332 |
| Anti-CD31 (PECAM) | Mouse | 1:100 | Biomedica | 303102 |
| Anti-Cx43 (Connexin 43,Gja1) | Rabbit | 1:400 | Sigma Aldrich | C6219-100UL |
| Anti-GATA4 | Rabbit | 1:400 | Cell signalling | 36966S |
| Anti-MESP1 | Rabbit | 1:200 | Abcam | ab129387 |
| Anti-MYL2 (α-MLC2) | Rabbit | 1:200 | Abcam | ab79935 |
| Anti-MYL7 (MLC2a) | Mouse | 1:200 | Santa Cruz | sc-365255 |
| Anti-NFAT2 (NFATc1) | Rabbit | 1:100 | Cell Signalling | 8032S |
| Anti-SERCA2A (ATP2A2) | Mouse | 1:100 | Novus Biologicals | NB300-518 |
| Anti-SOX2 | Rabbit | 1:400 | Sigma Aldrich | HPA015774-100UL |
| Anti-TE - 7 | Mouse | 1:100 | Sigma Aldrich | CBL271 |
| Anti-TNNT2 | Mouse | 1:200 | Thermo Fisher | MA5-12960 |
| Anti-TNNT2 | Rabbit | 1:3000 | Sigma Aldrich | HPA015774 |
| ***IF- Secondary Antibodies*** | | | | |
| **Name** | | **Dilution** | **Company** | **Catalog no** |
| Alexa Fluor 488 Donkey anti-Rabbit IgG (H+L) | | 1:500 | Thermo Fisher Scientific | A-21206 |
| Alexa Fluor 488 Donkey anti-Mouse IgG (H+L) | | 1:500 | Thermo Fisher Scientific | A-21202 |
| Alexa Fluor 555 Donkey anti-Rabbit IgG (H+L) | | 1:500 | Thermo Fisher Scientific | A-31572 |
| Alexa Fluor 555 Donkey anti- Mouse IgG (H+L) | | 1:500 | Thermo Fisher Scientific | A-31570 |
| Alexa Fluor 546 Donkey anti-Sheep IgG (H+L) | | 1:500 | Thermo Fisher Scientific | A-21098 |
| Alexa Fluor 647 Donkey anti-Goat IgG (H+L) | | 1:500 | Thermo Fisher Scientific | A-21447 |
| Alexa Fluor 647 Goat anti-Mouse IgG (H+L) | | 1:500 | Thermo Fisher Scientific | A-21235 |
| **FACS** | | | | |
| **Name** | | **Dilution** | **Company** | **Catalog no** |
| Anti Cardiac Troponin T-FITC Clone REA400 (REAfinity™) | | 1:50 | Miltenyi Biotec | 130-119-575 |
| Anti-Hu CD90 APC | | 1:10 | Exbio | 1A-652-T100 |
