## Supplementary Figures for "Generation and Maturation of Human iPSC-derived Cardiac Organoids in Long Term Culture"

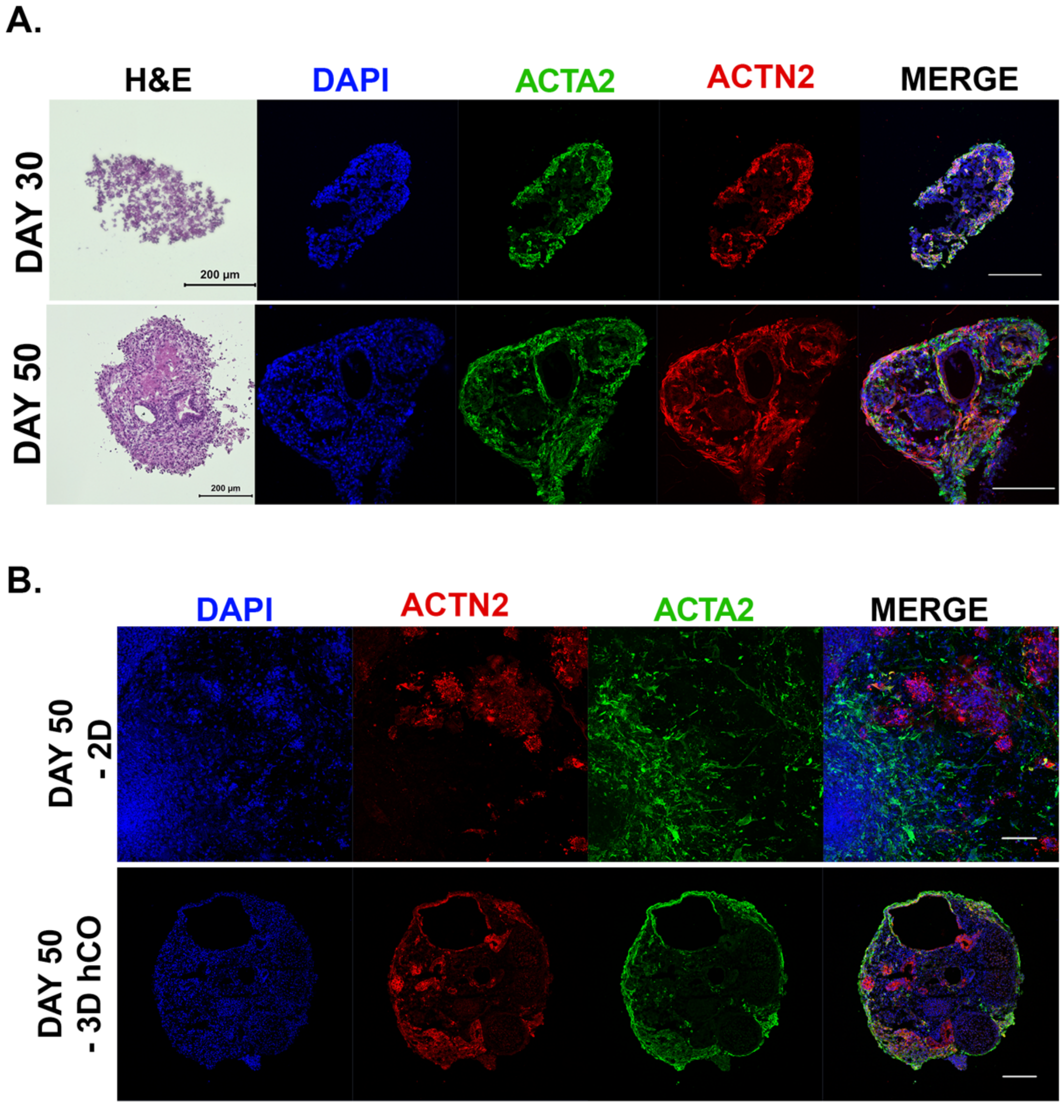


**Figure SI-1 - 3D hCOs recapitulate advanced morphology and cellular heterogeneity in longer culture times, compared to 2D monolayer culture**

**a.** H&E staining, and immunofluorescence analysis showing the increasing histological complexity of 3D hCO sections on day 30 vs day 50. **b.** IF analysis of 2D monolayer culture (tile-scan of one well), vs. 3D hCO sections on day 50 of culture. For both IF analyses, markers for cardiomyocytes (ACTN2), smooth muscle/fibroblastic cells (ACTA2), shown, counterstained with DAPI (blue). Scale bars = 200µm

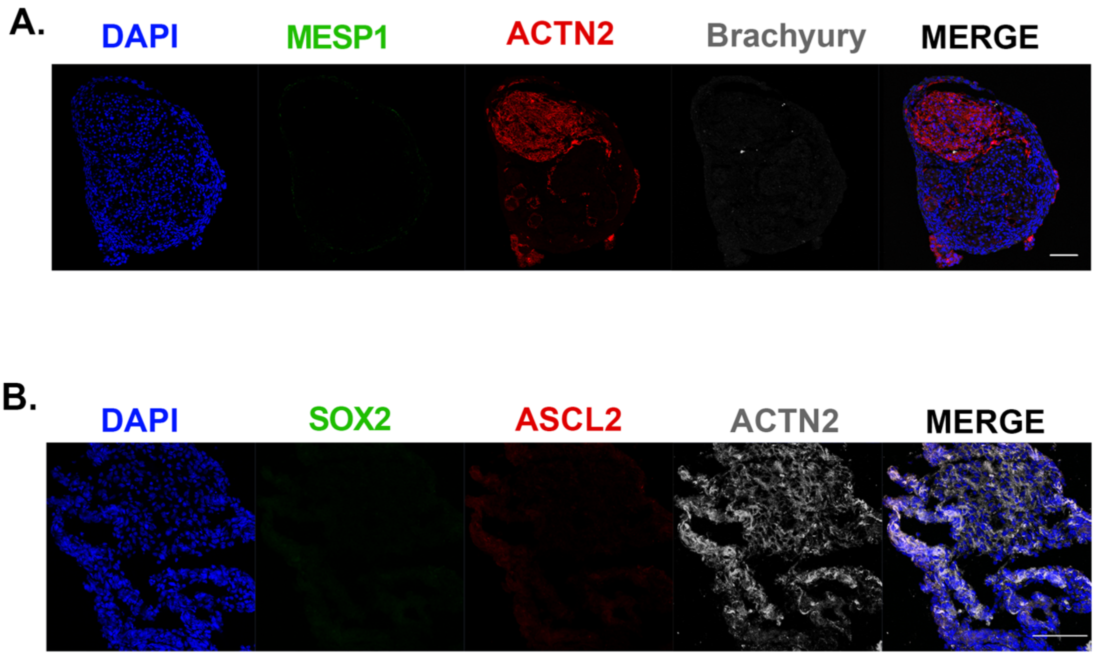


**Figure SI-2 – Long term hCOs do not show markers for undifferentiated cells or co-emergence of gut tissue.**

**a.** IF staining of hCO sections for undifferentiated cell markers including MESP1 and Brachyury, compared to differentiated cardiomyocytes (ACTN2) on day 50. Counterstaining with DAPI (blue). Scale bar = 100µm. **b.** IF staining of hCO sections for undifferentiated cell & embryonic gut (SOX2), and adult gut (ASCL2) tissue markers, compared to differentiated cardiomyocytes (ACTN2) on day 50. Counterstaining with DAPI (blue). Scale bar = 100µm

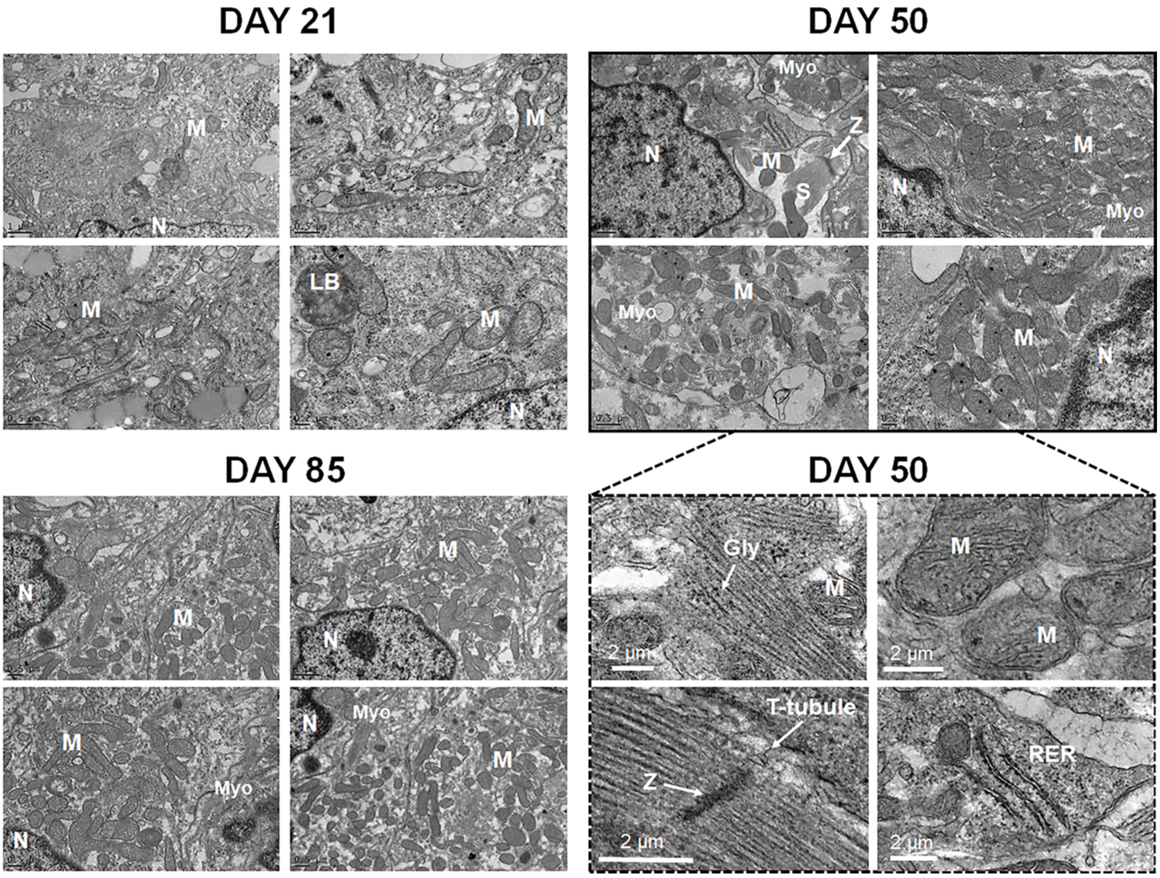


**Figure SI-3 – Ultrastructural analysis of hCOs indicates enhanced cardiomyocyte and metabolic maturation in 3D environment over time.**

TEM images of hCOs increasing ultrastructural organization and structural & metabolic maturity over time, from day 21 to day 85: Depicted: Cardiomyocyte myofibers (Myo), Sarcomeres (S), Mitochondria (M) Z-band (Z), T-tubules, Nuclei (N), Rough endoplasmic reticulum (RER), Glycogen granules (Gly), Lamellar bodies (LB). Scale bars are 1 µm and 0.5 µm for day 21, 0.5 µm for day 85, 0.5 µm for day 50 and 2 µm for magnified images of day 50.

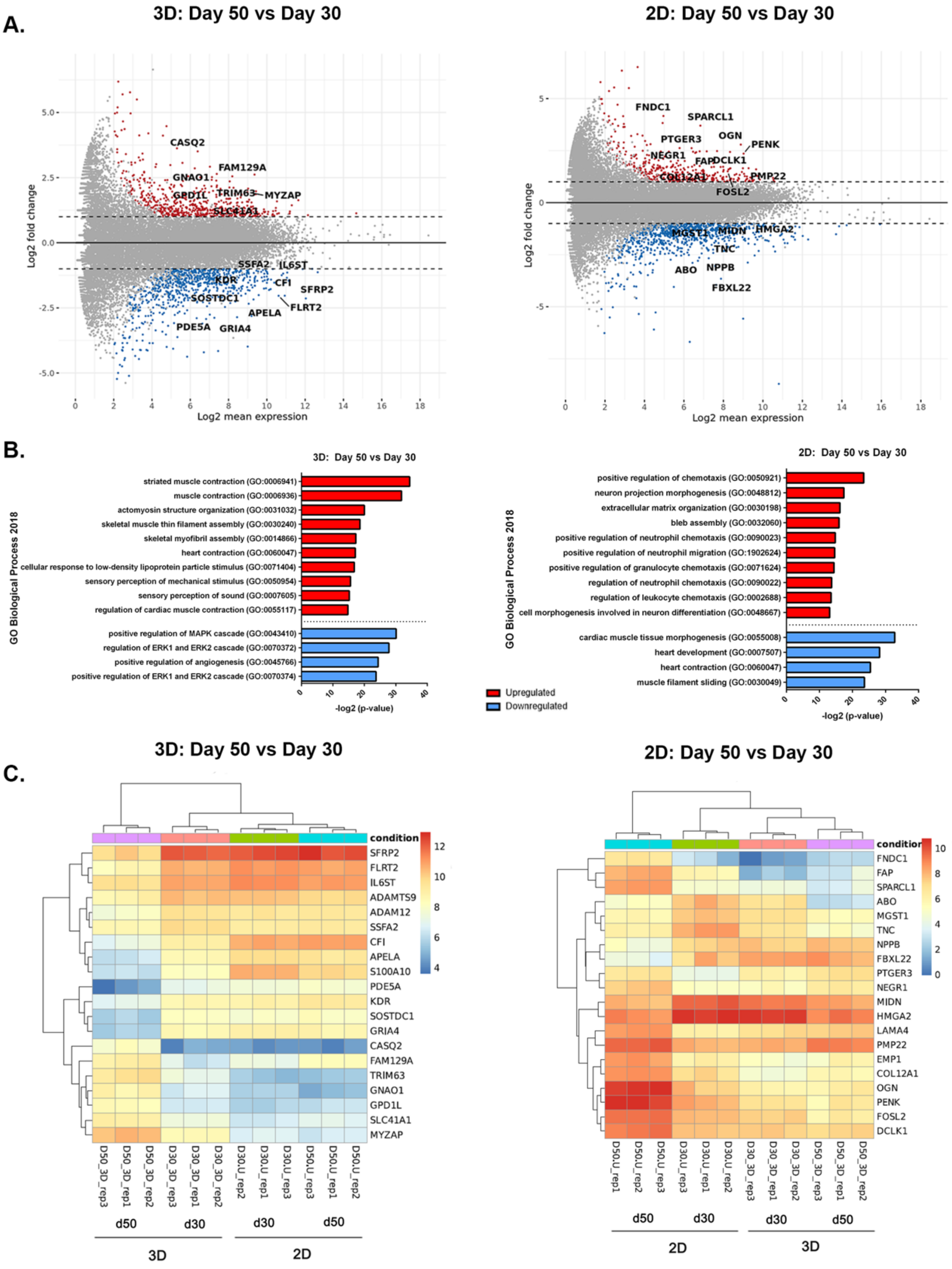


**Figure SI-4 – 3D l hCO cultures induce improved cardiac specification and cardiomyocyte maturation at a transcriptomic level with respect to culture dimensionality and time.**

**a.** MA plot of the differentially regulated genes between day 30 and day 50 for 3D hCOs (left) and 2D monolayer culture (right). **b.** Graph representing the -log2value adjusted p-value of significantly upregulated (red) or downregulated (blue) GO Biological process categories when comparing between day 30 and day 50 of culture for 3D hCOs (left) and 2D monolayer culture (right). **c.** Heatmap representing the log2fold change for the top 20 differentially regulated genes between day 30 and day 50 of culture for 3D hCOs (left) and 2D monolayer culture (right).
