## Supplementary Methods for "Generation and Maturation of Human iPSC-derived Cardiac Organoids in Long Term Culture"

### Histology and Immunofluorescence (IF)

After organoid fixation and cryosectioning, to assess the overall histology, the sections were stained with hematoxylin and eosin (H&E) (Sigma Aldrich), and imaged under slide scanner Zeiss Axio Scan Z1 microscope using the bright-field mode.

In addition to 3D hCO cryosections, 2D monolayer cultures were used as controls. As 2D monolayer controls were grown adherently on Matrigel-coated plates, for the IF procedure, the samples were fixed with 4% PFA for 15 minutes, and rest of the IF precedures were performed as previously described in the main text. The resulting samples were stored in PBS with 0.01% Sodium Azide (VWR) at 4℃, and protected from light until confocal image analysis.

See Supplementary table 3 for full list of antibodies used in the study.

### RNA-sequencing and Differential Expression analysis (DE analysis)

#### *Sequencing library preparation*

500 nanograms of total RNA per sample was used as an input for library preparation using QuantSeq FWD 3'mRNA Library Prep Kit (Lexogen). Briefly, RNA was transcribed into cDNA using oligodT primer (FS1) at half volume compared to the manufacturer's instructions to minimize the off-target products. Following the first strand synthesis, RNA removal and second strand synthesis were performed using the UMI Second Strand Synthesis Mix (USS) containing Unique Molecular Identifiers (UMIs) that allow detection and removal of PCR duplicates. Finally, sequencing libraries were created by PCR with i5 Unique Dual Indexing Add-on Kit for Illumina (Lexogen). The quality and quantity of libraries was determined using Fragment Analyzer by DNF-474 High Sensitivity NGS Fragment Analysis Kit (Agilent Technologies) and QuantiFluor dsDNA System (Promega). Final library pool was sequenced on Illumina NextSeq 500 with 75 bp single-ends, producing about 10 million reads per library.

#### *Data analysis*

High-throughput RNA-Seq data were prepared using Lexogen QuantSeq 3' mRNA-Seq Library Prep Kit FWD for Illumina with polyA selection and sequenced on Illumina NextSeq 500 sequencer (run length 1x75 nt). Bcl files were converted to Fastq format using bcl2fastq v. 2.20.0.422 Illumina software for basecalling. 6-nt long UMIs were extracted and subseqeuntly used for deduplication of aligned reads by UMI-tools v. 1.1.1. (1). As a next step 6-nt long barcode sequence related to Lexogen QuantSeq Library Prep Kit were trimmed using seqtk 1.3-r106 (2). Quality check of raw single-end fastq reads was carried out by FastQC v0.11.9 (3). The adapters and quality trimming of raw fastq reads was performed using Trimmomatic v0.36 (4) with settings CROP:250 LEADING:3 TRAILING:3 SLIDINGWINDOW:4:5 MINLEN:35 and adaptor seqeunce ILLUMINACLIP:AGATCGGAAGAGCACACGTC. Trimmed RNA-Seq reads were mapped against the human genome (hg38) and Ensembl GRCh38 v.94 annotation using STAR v2.7.3a (5) as splice-aware short read aligner and default parameters except --outFilterMismatchNoverLmax 0.1 and --twopassMode Basic. Quality control after alignment concerning the number and percentage of uniquely and multi-mapped reads, rRNA contamination, mapped regions, read coverage distribution, strand specificity, gene biotypes and PCR duplication was performed using several tools namely RSeQC v2.6.2 (6), Picard toolkit v2.18.27 (7) and Qualimap v.2.2.2 (8) and BioBloom tools v 2.3.4-6-g433f (9).

The differential gene expression analysis was calculated based on the gene counts produced using featureCounts tool v1.6.3 (10) with settings -s 2 -T 10 -F GTF -Q 0 -d 1 -D 25000 and using Bioconductor package DESeq2 v1.20.0 (11). Data generated by DESeq2 with independent filtering were selected for the differential gene expression analysis to avoid potential false positive results. Genes were considered as differentially expressed based on a cut-off of adjusted p-value <= 0.05 and log2(fold-change) ≥1 or ≤-1. Clustered heatmaps were generated from selected top differentially regulated genes using R package pheatmap v1.0.10 (12). Volcano plots were produced using ggplot v3.3.3 package (13) and MA plots were genereted using ggpubr v0.4.0 package (14).

See Supplementary table 1 for log2changes and p-values for the different comparisons, and Supplementary table 2 for GO BP clustering analysis for the differentially regulated genes at the different comparisons.
